# TomatoPGFM: A graph-conditioned foundation model for tomato pangenomes

**DOI:** 10.64898/2026.08.31.748176

**Authors:** Jinlei Han, Tuerhong Yushan, Juan Wang, Haitao Yang, Jiantao Zhao, Fangling Jiang, Chunping Jia, Tao Yang, Baike Wang, Changwei Zhang, Qinghui Yu

## Abstract

Most genomic foundation models are pretrained on independent linear assemblies and therefore do not explicitly represent population-level segment sharing or local graph connectivity. We developed TomatoPGFM, a graph-conditioned model pretrained on 54.65 Gb of sequence from 66 tomato (*Solanum* spp.) accessions. Sequence tokens were conditioned on pangenome node attributes and local adjacency, and the model was optimised using masked language modelling and graph-feature reconstruction. To evaluate model responses to graph-conditioned input, we compared aligned, shuffled and disabled graph inputs in 25, 000 windows from the training panel. Sequence-aligned graph input produced lower masked language modelling loss than graph-off at all five curriculum stages in both training-panel strata, while the shuffled perturbation generally yielded intermediate losses. We then assessed sequence-only transfer in *Solanum sitiens* LA1974 and *S. lycopersicum* MicroTom, neither of which was used for graph construction or pretraining. Frozen-probe AUROC values for gene-versus-intergenic and coding-sequence-versus-intergenic classification ranged from 0.8489 to 0.9593. TomatoPGFM produced higher AUROC point estimates than DNABERT-2 in all four comparisons. Enabling the zero-feature GraphAdapter pathway with adjacency messaging disabled changed throughput by less than 1% at 512–2, 048 positions under the tested configuration. Together, these results show that TomatoPGFM responds consistently to sequence-aligned pangenome context in training-panel sequences and provides informative sequence representations for genic-region classification in accessions excluded from graph construction and pretraining.

## Introduction

DNA sequence models can learn representations that support genome annotation, regulatory prediction and variant-effect estimation. Supervised long-context models have shown that distal sequence can improve regulatory prediction (Avsec et al., 2021), whereas self-supervised models based on masked-token or autoregressive objectives have extended representation learning across taxa and sequence scales (Ji et al., 2021; Nguyen et al., 2023; Zhou et al., 2024; Schiff et al., 2024; Dalla-Torre et al., 2025). Plant-focused and plant-inclusive models have been evaluated for variant effects, regulatory annotations, gene structure and cross-species transfer in *Arabidopsis*, crops and diverse angiosperms (Benegas et al., 2023; Mendoza-Revilla et al., 2024; Liu et al., 2025; Zhai et al., 2025; Li et al., 2026). However, benchmark performance depends strongly on the pretraining corpus, architecture, tokenisation, representation layer and pooling strategy (Feng et al., 2025). Species-focused models have also begun to use population-scale assemblies as pretraining corpora (Qian et al., 2026; Yang et al., 2026).

Most genomic foundation models nevertheless treat each assembly as an independent linear sequence. A single reference cannot capture all presence/absence and structural variation within a plant species, as shown in crop pangenomes (Golicz et al., 2016; Bayer et al., 2020). A pangenome represents shared, variable and accession-specific sequence, while a graph pangenome additionally encodes segment connectivity and accession paths (Computational Pan-Genomics Consortium, 2018; Eizenga et al., 2020; Hickey et al., 2024). Plant graph frameworks use path information to store haplotypes and support imputation or population analysis (Bradbury et al., 2022). In tomato, pangenome studies have identified genes absent from the reference assembly, structural variants associated with gene expression, and alleles linked to fruit quality, stress responses and breeding traits (Gao et al., 2019; Alonge et al., 2020; Zhou et al., 2022; Li et al., 2023; Shi et al., 2026). Graph nodes can therefore provide both the training-panel prevalence of a segment and its local genomic neighbourhood, neither of which is explicit when assemblies are supplied only as separate sequence documents.

How such graph structure should be incorporated into a genomic foundation model, and how its contribution should be tested, remain unresolved. OneGenome-Rice and OryzaG3 broaden the pretraining corpus by using multiple rice assemblies as sequence documents (Qian et al., 2026; Yang et al., 2026), whereas DeepGene serialises sequence and positional information derived from a pangenome graph (Zhang et al., 2025). An alternative is to retain nucleotide sequence as the primary input while conditioning token representations directly on graph attributes and local adjacency. This design raises two questions: whether sequence-aligned graph context contributes to sequence modelling for accessions represented in the graph, and whether the sequence encoder remains useful when graph coordinates are unavailable.

Here, we developed TomatoPGFM from a 66-accession tomato pangenome by combining graph-node attributes and adjacency-based message passing with a hybrid long-context sequence backbone. We compared sequence-matched, shuffled and disabled graph inputs in windows from the training panel to test whether masked-token prediction depended on sequence-graph correspondence. We then evaluated sequence-only transfer to *S. sitiens* LA1974 and *S. lycopersicum* MicroTom, both excluded from graph construction and pretraining, using chromosome-separated gene-versus-intergenic and coding-sequence-versus-intergenic classification with frozen probes and low-rank adaptation.

## Results

### A 66-accession pangenome captures segment prevalence and local connectivity

The training panel comprised 66 accessions spanning cultivated *Solanum lycopersicum* sensu lato and seven wild relatives. Their chromosome-scale assemblies totalled 54.65 Gb and contained 2, 269, 697 annotated genes. *Solanum sitiens* LA1974 and the *S. lycopersicum* cultivar MicroTom were reserved for downstream evaluation, and neither accession contributed to graph construction, tokeniser construction, pretraining-shard generation or model pretraining. With SL6.0 Heinz 1706 as the backbone, minigraph generated 3, 203, 440 segments and 4, 450, 064 links. The graph contained 2.598 Gb of sequence. Segment length had a median of 114 bp and a 95th percentile of 3, 978 bp, while node degree had a median of 2 and a 95th percentile of 4 (Figure 1; Table S1).

**Figure 1.**
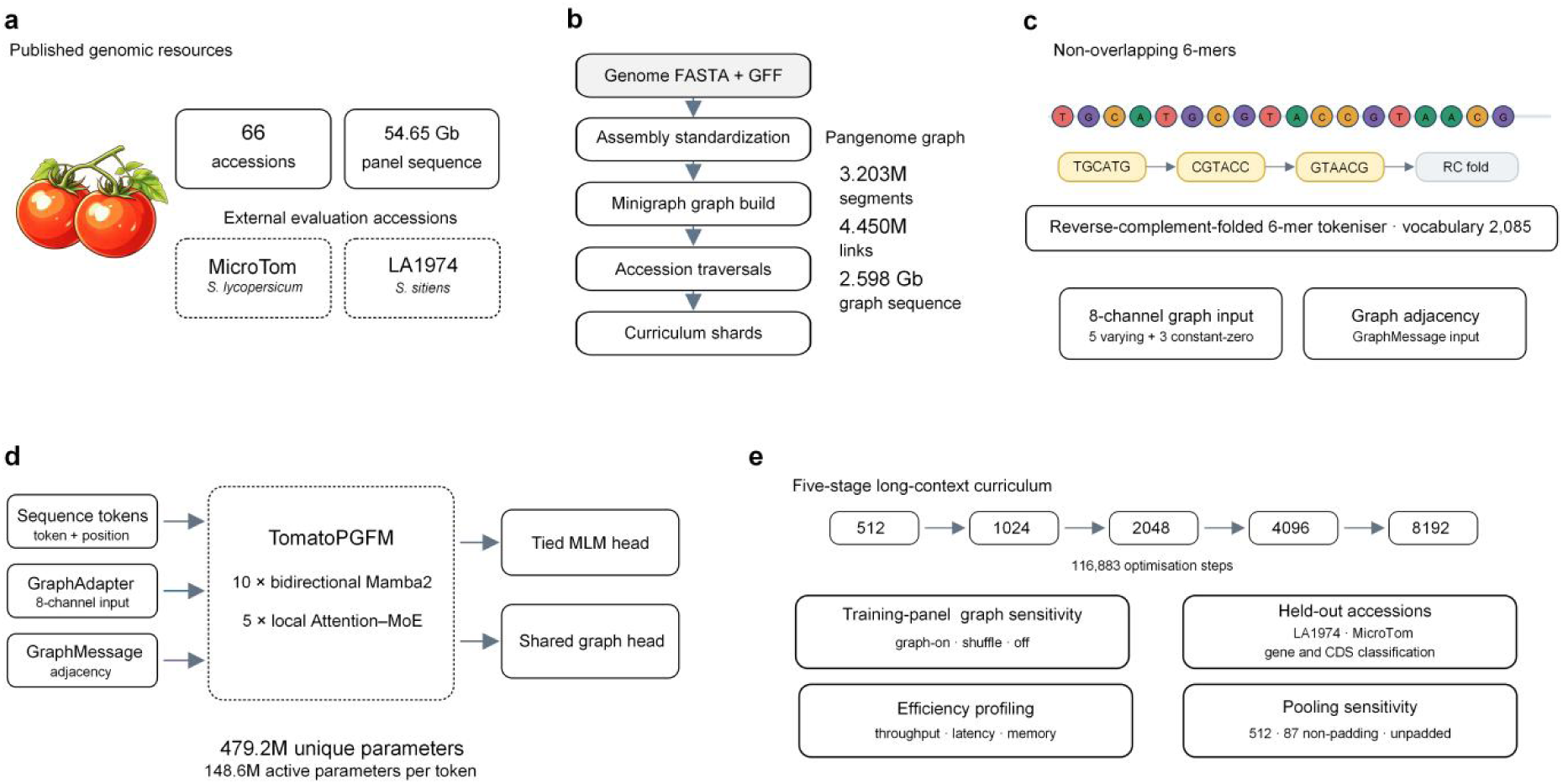
TomatoPGFM development and evaluation workflow. (a) The 66-accession, 54.65 Gb pretraining panel, with LA1974 and MicroTom excluded from graph construction and pretraining. (b) Assembly and annotation standardisation, minigraph construction, calculation of traversal-derived features and generation of curriculum shards, with graph summary statistics. (c) Reverse-complement-folded 6-mer tokenisation, an eight-channel graph interface containing five variable segment-level channels, including sequence-, topology- and traversal-derived quantities, plus three constant-zero channels, and within-window adjacency. (d) TomatoPGFM architecture, comprising GraphAdapter, GraphMessage, 10 Mamba2 blocks, five local-attention mixture-of-experts blocks, a tied masked language modelling head and a shared graph-reconstruction head. (e) Pretraining and evaluation analyses, including graph-input sensitivity in the training panel, gene and CDS classification in held-out accessions, inference efficiency and pooling sensitivity.

Accession paths were used to count how many training-panel genomes traversed each segment. Among the 3, 178, 265 segments assigned to at least one path, 170, 034 were traversed by all 66 accessions. In contrast, 1, 315, 249 segments were traversed by one to six accessions, including 506, 198 found on a single accession path. The graph therefore represented both broadly shared and sparsely distributed sequence segments.

### TomatoPGFM conditions long-context sequence representations on node attributes and adjacency

TomatoPGFM represents sequence as non-overlapping reverse-complement-folded 6-mers and incorporates pangenome information through two gated residual pathways. GraphAdapter projects the eight-channel token-aligned segment interface through a gated residual pathway, whereas GraphMessage adds mean-aggregated messages from incoming within-window graph neighbours. The detailed channel definitions and their provenance are provided in the Methods. GraphMessage mean-aggregates hidden states from incoming graph neighbours and injects the resulting messages through a separate gated residual projection. The two modules feed a 15-block hybrid backbone comprising 10 bidirectional Mamba2 blocks and five local-attention mixture-of-experts blocks. The model contained 479, 195, 678 unique parameters, of which approximately 148, 625, 438 were active per token (Figure 1).

Pretraining combined masked language modelling with reconstruction of the eight-channel graph-feature target at all feature-valid positions and, separately, at masked-token positions. The first five target channels varied across production examples, whereas the remaining three channels were constant zero. The two graph losses shared an eight-dimensional output head and each was weighted at one-quarter of the masked language modelling objective. A five-stage curriculum increased model length from 512 to 8, 192 positions over 116, 883 optimisation steps. Training was completed in 46.09 h on four 80-GB NVIDIA A100 GPUs (Figure 2).

**Figure 2.**
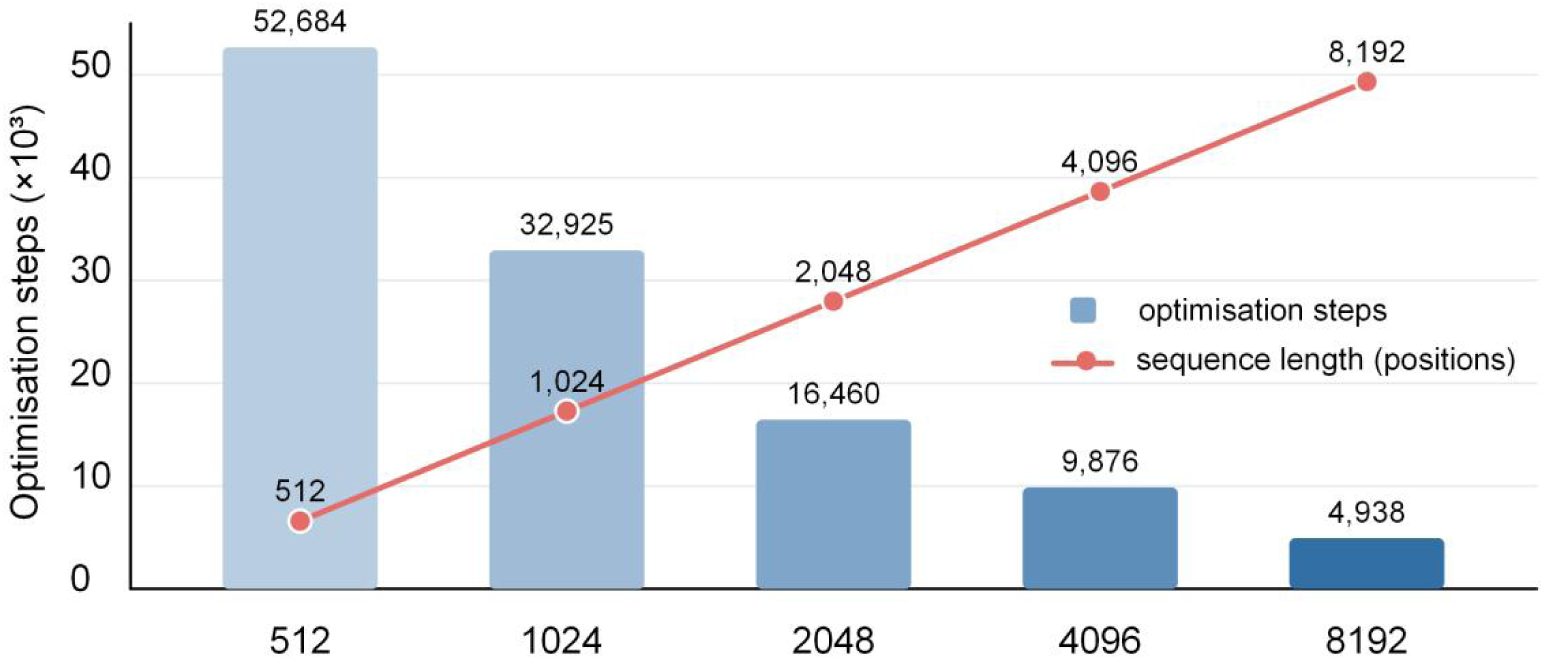
Five-stage pretraining curriculum of TomatoPGFM. The training sequence length progressively increased from 512 to 8, 192 model positions, while the corresponding number of optimisation steps decreased from 52, 684 to 4, 938. The curriculum used 52, 684, 32, 925, 16, 460, 9, 876 and 4, 938 optimisation steps at sequence lengths of 512, 1, 024, 2, 048, 4, 096 and 8, 192 positions, respectively. Bars show optimisation steps, with exact step counts labelled above each bar. The red line traces the increase in sequence length, with the corresponding number of model positions labelled at each point.

### TomatoPGFM responds to sequence-aligned graph inputs across pretraining stages

To evaluate model responses to graph input, we compared graph-on, graph-shuffle and graph-off inference at the end of each curriculum stage using identical batches. Graph-on retained sequence-matched node features and adjacency. Graph-shuffle permuted node features along the sequence axis within each example and destination indices across the flattened batch; this preserved the feature-row multiset, total edge count and marginal source out-degree and destination in-degree counts, but could create cross-example links and changed source– destination pairings. Graph-off disabled both conditioning pathways. The analysis included 25, 000 windows from the 66-accession training panel: 21, 750 wild-panel windows and 3, 250 cultivated-panel windows. Graph-on yielded lower masked language modelling loss than graph-off at all five stages in both strata. In the wild-panel stratum, the graph-on minus graph-off difference was −0.1265 nats at 512 positions (95% paired-bootstrap interval, −0.1282 to −0.1248) and −0.0833 nats at 8, 192 positions (−0.0846 to −0.0820). The shuffled perturbation generally yielded intermediate losses, demonstrating consistent sensitivity to the supplied graph-conditioned input across pretraining stages (Figure 3). We interpret this result as a joint graph-input sensitivity test, not as an isolated topology-specific intervention.

**Figure 3.**
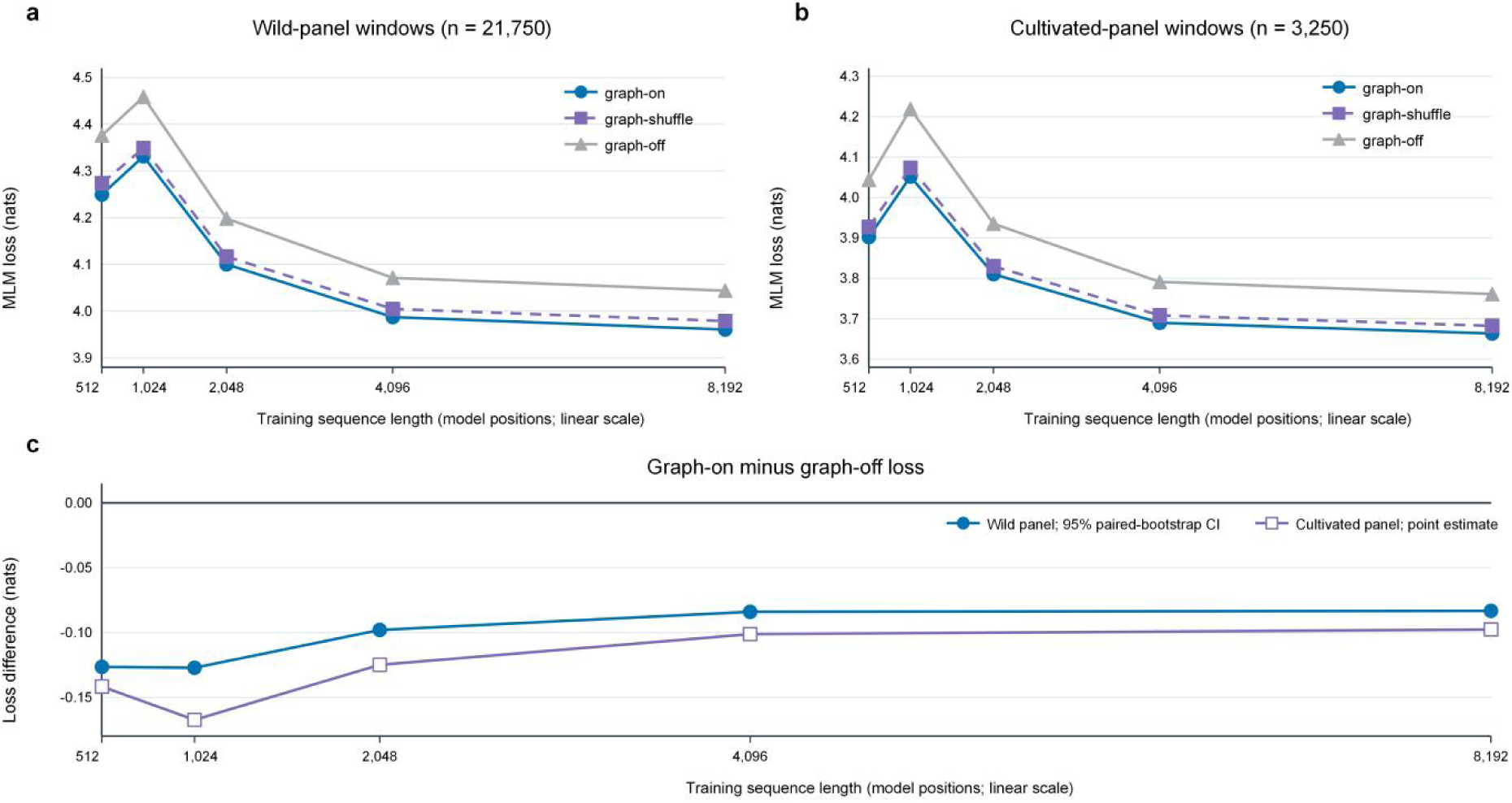
Sensitivity to sequence-aligned graph input in training-panel windows. (a, b) Masked language modelling loss in graph-on, graph-shuffle and graph-off modes at the five curriculum checkpoints for wild-panel windows (n = 21, 750) and cultivated-panel windows (n = 3, 250). (c) Difference between graph-on and graph-off loss. Wild-panel estimates show 95% paired-bootstrap intervals from 1, 000 resamples; cultivated-panel values are point estimates. Negative values indicate lower loss with sequence-aligned graph input. The graph-shuffle condition is a joint input perturbation; its destination permutation can create cross-example links and is not a topology-preserving null.

### Sequence-only representations distinguish genic from intergenic sequence in held-out accessions

To assess transfer independently of graph mapping, we generated balanced gene-versus-intergenic and coding-sequence (CDS)-versus-intergenic datasets for LA1974 and MicroTom. Positive windows were centred within annotated gene or CDS intervals, negative windows were entirely intergenic, and windows crossing annotation boundaries were excluded. Chromosomes 03 and 05 were reserved for final testing, thereby separating classifier fitting and model selection from the test chromosomes. TomatoPGFM was evaluated without graph input. Each 512-bp window was represented by the mean hidden state across its 87 non-padding positions, including the two special tokens, whereas DNABERT-2 and PlantDNAMamba were processed with their native tokenisers and attention-mask-aware pooling. StandardScaler and class-balanced logistic regression were fitted together within each training fold. Three-fold stratified cross-validation selected the regularisation parameter C from seven prespecified values; the selected pipeline was refitted on all training-chromosome samples and evaluated once on chromosomes 03 and 05. Uncertainty was estimated from 1, 000 class-stratified bootstrap resamples of the fixed test predictions.

PlantDNAMamba had the highest AUROC point estimate in each of the four accession-by-task comparisons, followed by TomatoPGFM and DNABERT-2. TomatoPGFM achieved AUROC values of 0.8638 (95% bootstrap interval, 0.8513–0.8760) for LA1974 gene classification and 0.9592 (0.9473–0.9710) for LA1974 CDS classification. The corresponding MicroTom values were 0.8489 (0.8354–0.8616) and 0.9593 (0.9482–0.9695). TomatoPGFM produced higher AUROC point estimates than DNABERT-2 by margins of 0.0387–0.0531 but remained below PlantDNAMamba. Its sequence-only representations therefore distinguished gene and CDS windows from intergenic windows in both accessions excluded from pretraining (Figure 4; Table 1).

**Figure 4.**
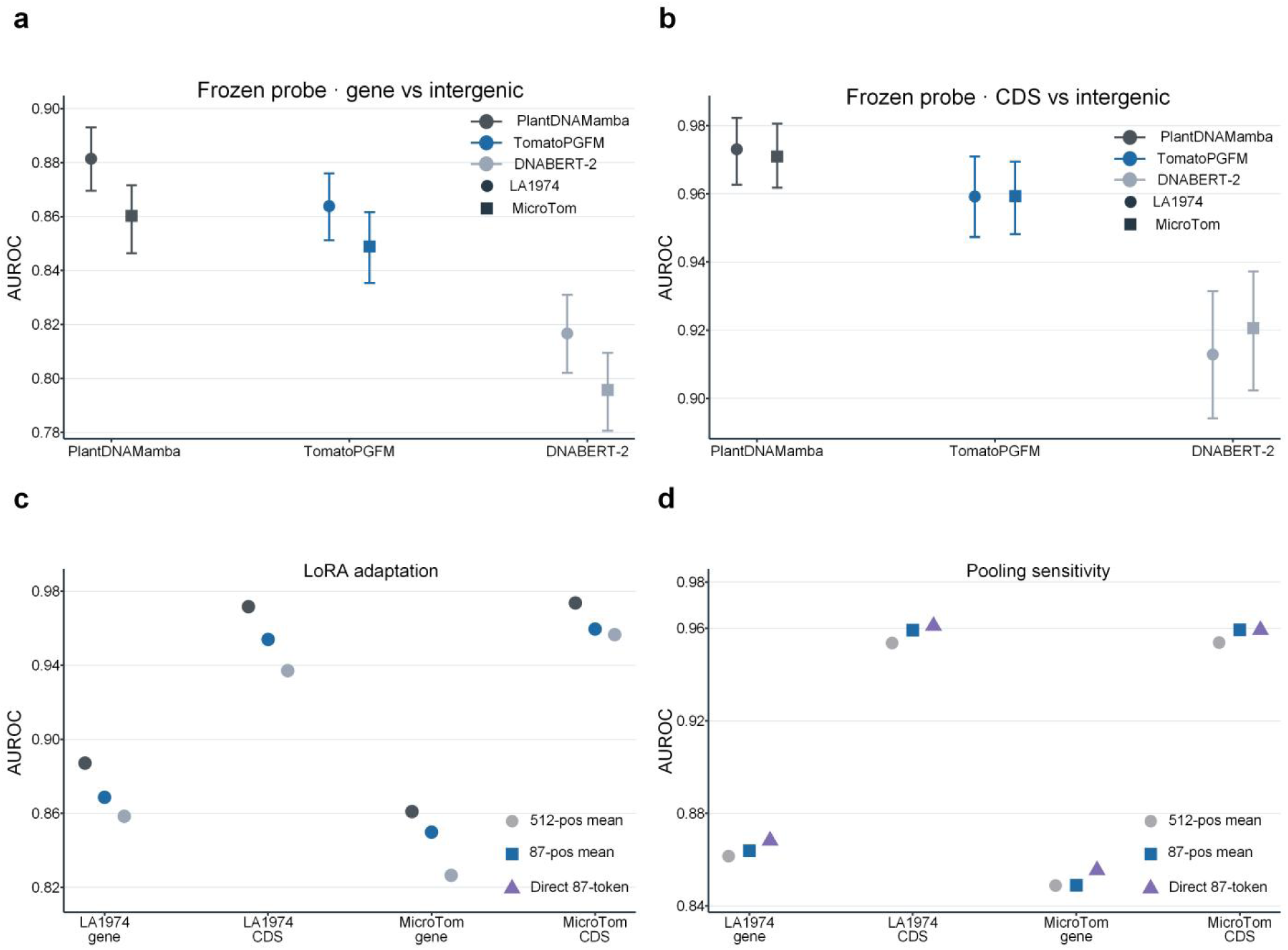
Sequence-only evaluation in held-out tomato accessions. (a, b) Frozen-probe AUROC values for gene-versus-intergenic and CDS-versus-intergenic classification in LA1974 and MicroTom. Error bars show 95% class-stratified bootstrap intervals from 1, 000 resamples. Frozen-probe test sets contained 3, 012 and 3, 836 windows for gene classification and 808 and 910 windows for CDS classification in LA1974 and MicroTom, respectively. (c) LoRA AUROC point estimates. Gene test sets contained 3, 012 LA1974 and 3, 836 MicroTom windows; CDS test sets contained 767 LA1974 and 885 MicroTom windows. (d) TomatoPGFM AUROC after pooling all 512 executed positions, pooling the 87 non-padding positions, or directly executing the unpadded 87-token input.

**Table 1.**
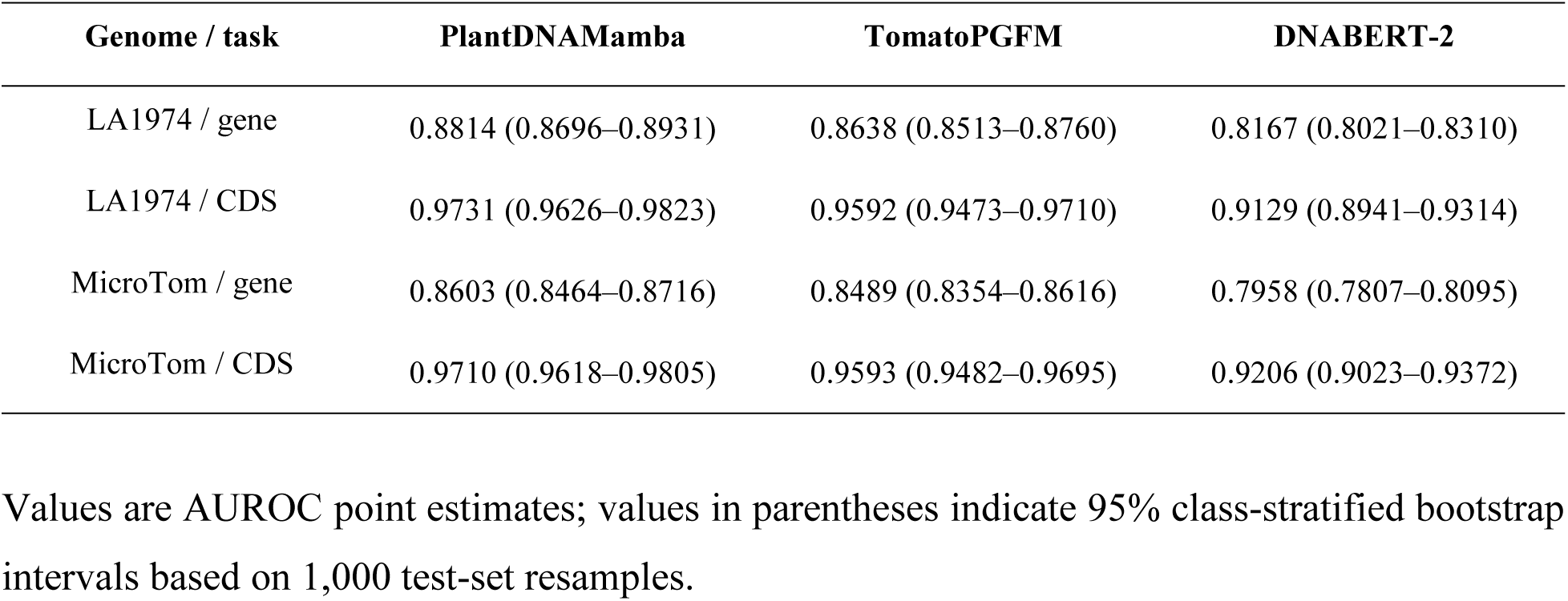
Frozen-probe AUROC values for sequence-only classification in held-out accessions.

| Genome / task | PlantDNAMamba | TomatoPGFM | DNABERT-2 |
| --- | --- | --- | --- |
| LA1974 / gene | 0.8814 (0.8696–0.8931) | 0.8638 (0.8513–0.8760) | 0.8167 (0.8021–0.8310) |
| LA1974 / CDS | 0.9731 (0.9626–0.9823) | 0.9592 (0.9473–0.9710) | 0.9129 (0.8941–0.9314) |
| MicroTom / gene | 0.8603 (0.8464–0.8716) | 0.8489 (0.8354–0.8616) | 0.7958 (0.7807–0.8095) |
| MicroTom / CDS | 0.9710 (0.9618–0.9805) | 0.9593 (0.9482–0.9695) | 0.9206 (0.9023–0.9372) |

Values are AUROC point estimates; values in parentheses indicate 95% class-stratified bootstrap intervals based on 1,000 test-set resamples.

TomatoPGFM performance was similar across the three pooling procedures. Relative to the primary non-padding mean, pooling all 512 executed positions changed AUROC by −0.0023, −0.0056, −0.0001 and −0.0055 across the four evaluations. Direct execution of the unpadded 87-token input changed AUROC by +0.0044, +0.0019, +0.0065 and −0.0001. All absolute differences were at most 0.0065, and the model ranking was unchanged (Figure 4d).

### Low-rank adaptation retains the frozen-probe model ranking

We next applied low-rank adaptation (LoRA) to the same held-out-accession tasks. Architecture-matched projection modules were adapted with rank 8 and scaling factor 16, yielding 1, 193, 218 trainable parameters for TomatoPGFM, 456, 194 for DNABERT-2 and 1, 353, 218 for PlantDNAMamba. A random 10% subset of the training-chromosome samples was used for validation and early stopping. Gene test sets were unchanged from the frozen-probe analysis, whereas the balanced CDS test sets contained 767 LA1974 windows and 885 MicroTom windows. LoRA retained the same model ranking in all four comparisons. TomatoPGFM achieved AUROC values of 0.8687 and 0.9540 for LA1974 gene and CDS classification, respectively, compared with 0.8584 and 0.9371 for DNABERT-2 and 0.8872 and 0.9717 for PlantDNAMamba. For MicroTom, TomatoPGFM achieved 0.8499 and 0.9596, compared with 0.8265 and 0.9566 for DNABERT-2 and 0.8610 and 0.9737 for PlantDNAMamba. TomatoPGFM point estimates were 0.0030–0.0234 higher than those of DNABERT-2 and 0.0111–0.0185 lower than those of PlantDNAMamba (Figure 4c).

### Zero-feature GraphAdapter activation has minimal measured inference overhead

Enabling the zero-feature GraphAdapter pathway with adjacency messaging disabled added little computational overhead under the tested configuration. On one 80-GB NVIDIA A100 GPU, with a batch size of 16 and FP32 inference, zero-feature adapter-on and graph-off throughput differed by less than 1% at 512, 1, 024 and 2, 048 model positions. At 2, 048 positions, zero-feature adapter-on and graph-off processed 23.05 and 23.16 sequences per second, respectively. Median latency was 93.16 ms for zero-feature adapter-on and 92.12 ms for graph-off, and peak memory was 3, 389.9 MB in both modes. At 512 positions, peak memory was 2, 370.1 MB with the zero-feature GraphAdapter pathway enabled and 2, 226.1 MB in graph-off mode. For a 512-bp window, TomatoPGFM executed 512 model positions but retained 87 non-padding tokens: 85 sequence-derived 6-mers and two special tokens. In graph-off mode, 78.41 sequences per second corresponded to 40, 147 model positions, 6, 822 non-padding tokens, 6, 665 6-mer tokens and 40, 146 input base pairs per second (Figure 5).

**Figure 5.**
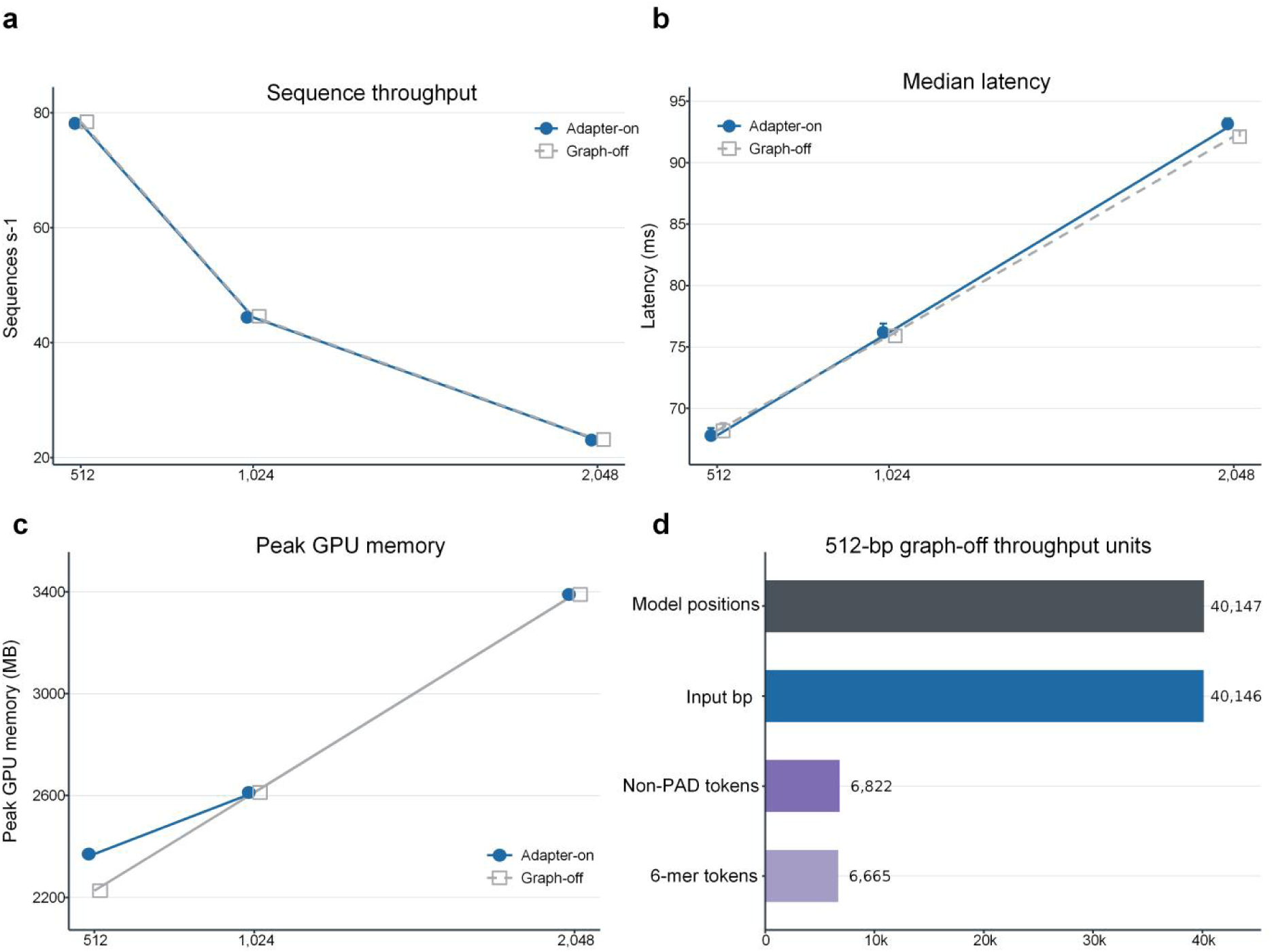
TomatoPGFM inference efficiency with the zero-feature GraphAdapter pathway enabled or disabled. (a) Sequence throughput in adapter-on and graph-off modes. (b) Median latency, with upper whiskers showing the recorded 90th percentile. (c) Peak GPU memory. (d) Throughput units for the 512-bp graph-off benchmark. The adapter-on condition used zero-valued graph features with edge_index = None, so adjacency-based GraphMessage aggregation was not executed. Measurements used one 80-GB NVIDIA A100 GPU, batch size 16 and FP32 inference.

## Discussion

TomatoPGFM was sensitive to supplied graph context when it was aligned with sequence, whereas its sequence-only representations transferred to two accessions excluded from graph construction and pretraining. In training-panel windows, sequence-aligned graph input produced lower masked language modelling loss than graph-off at every curriculum stage, while the batch-shuffled perturbation generally produced intermediate losses. These results demonstrate consistent sensitivity to the supplied graph-conditioned input across pretraining stages. The training-panel analysis and held-out-accession analysis therefore address complementary model properties. This analysis establishes sensitivity to the supplied graph-conditioned input during the MLM probe; it does not isolate the contribution of an individual feature group, topology component or auxiliary loss.

LA1974 and MicroTom provided a separate test of representation transfer because neither accession contributed to the training graph or pretraining corpus, and TomatoPGFM was evaluated as a sequence-only encoder. Its frozen representations had higher AUROC point estimates than DNABERT-2 in all four comparisons. The concordant ranking across frozen probing and parameter-efficient adaptation supports the conclusion that TomatoPGFM encodes sequence features that separate gene and CDS windows from intergenic sequence in held-out accessions. These comparisons do not isolate an effect of graph conditioning, because the three models differ in architecture, tokenisation, parameter count and pretraining corpus, all of which can affect genomic foundation-model benchmarks (Feng et al., 2025). Holding out chromosomes 03 and 05 separated classifier fitting from final testing and reduced local sequence overlap. The bootstrap intervals quantify uncertainty in the fixed frozen-probe predictions, whereas the LoRA analysis used one validation split and reports point estimates only.

TomatoPGFM differs from recent crop-specific models in how pangenome information enters the network. OneGenome-Rice and OryzaG3 use multiple rice assemblies as sequence corpora (Qian et al., 2026; Yang et al., 2026), whereas DeepGene serialises graph-derived sequence and positional information (Zhang et al., 2025). TomatoPGFM instead supplies node attributes and within-window adjacency as explicit conditioning signals while retaining nucleotide sequence as the primary input. This design allows sequence-only use when graph coordinates are unavailable and graph-conditioned use when they are available. Under the benchmarked hardware and batch settings, enabling the zero-feature GraphAdapter pathway with adjacency messaging disabled changed throughput by less than 1%, with the same recorded peak memory at 1, 024 and 2, 048 positions and a 144.0-MB increase at 512 positions. These efficiency measurements are specific to the tested implementation and conditions.

Tomato pangenomes have shown that presence/absence variation and structural variation can alter gene content, expression and agronomic traits (Gao et al., 2019; Alonge et al., 2020; Zhou et al., 2022; Shi et al., 2026). Tasks that depend directly on such intraspecific variation, including accession-dependent gene presence, structural-variant-associated expression and cis-regulatory activity, are therefore the most direct settings in which to test whether explicit pangenome context improves prediction. The gene- and CDS-versus-intergenic tasks used here do not answer that question, because TomatoPGFM was evaluated without graph input and no matched sequence-only pretraining control was included. Mapping held-out accessions to the pangenome would permit direct tests of graph-conditioned downstream inference, while additional accessions, chromosome partitions and training seeds would assess robustness. The present results establish two prerequisites for these experiments: sequence-aligned graph context affects masked-token prediction in training-panel sequences, and the sequence encoder transfers beyond the pretraining panel.

## Materials and methods

### Genome resources and pangenome construction

The pretraining panel comprised 66 accessions selected from three published genomic resources: 39 from Zhou et al. (2022), eight from Li et al. (2023), and 19 from Shi et al. (2026) after LA1974 was reserved for external evaluation. The panel totalled 54.65 Gb and contained 2, 269, 697 annotated genes. *Solanum sitiens* LA1974 (Molitor et al., 2021) and *S. lycopersicum* MicroTom (NCBI RefSeq GCF_036512215.1) were excluded from graph construction and pretraining and used only for downstream evaluation.

The pangenome graph was built with minigraph v0.21-r606 (Li et al., 2020) using SL6.0 Heinz 1706 as the backbone. Assemblies were added with -c -xggs and traversed with -cxasm, after which directed links were deduplicated and self-links removed. Carrier count was the number of training-panel paths containing each segment. GraphAdapter used eight token-aligned segment channels. Five varied in the production shards: sequence-derived segment length and GC fraction, topology-derived in-degree and out-degree, and traversal-derived accession carrier count. Segment-length, degree and carrier-count channels were encoded as ln(1+x); the remaining three channels were constant zero. The gene-overlap, repeat/transposable-element-overlap and reserved channels were constant zero. Token-validity masks and within-input adjacency were stored separately.

### Tokenisation, model architecture and pretraining

Sequences were encoded as non-overlapping reverse-complement-folded 6-mers using 2, 080 sequence tokens and five special tokens. Pretraining shards were generated without adding [CLS] and [SEP]. For downstream evaluation, each 512-bp window produced 85 complete 6-mers plus [CLS] and [SEP], yielding 87 non-padding positions. Sequence records with an N fraction greater than 0.30 were excluded during assembly standardisation, and graph segments with an N fraction greater than 0.50 were skipped during shard construction. Eligible training positions were masked using a window-ID-determined 15% Bernoulli procedure, with the first position excluded from masking.

In the production shard builder, each graph segment was tokenised independently into non-overlapping reverse-complement-folded (canonical) 6-mers at positions 0, 6, 12 and so on. Trailing sequence shorter than six bases was not emitted, and no 6-mer crossed a segment boundary. Every emitted token retained the source segment_idx and inherited the segment’s eight-dimensional feature vector. When a segment generated multiple tokens, all received the same feature vector; each directed segment link s → d contributed at most one token-level edge, defined by the first-token anchors of s and d when both segments were represented in the same window. For each source segment represented in a window, links to destination segments absent from that window were omitted and counted as outside-window projections. In the production shards, sequence-neighbour edges were not substituted for graph links. During collation, local edge indices were shifted by the example offset (i × max_len), where i denotes the example’s batch position and max_len the padded length of the batch; endpoints in padding positions were discarded. Consequently, the normal graph-on path did not create cross-example edges within a collated batch.

TomatoPGFM used a 768-dimensional, 15-block backbone comprising 10 bidirectional Mamba2 and five local-attention mixture-of-experts (MoE) blocks in a 2:1 pattern. Local attention used a 256-position window and 12 heads, and each MoE layer had 16 experts with top-2 routing (Shazeer et al., 2017; Fedus et al., 2022; Dao and Gu, 2024). GraphAdapter projected the eight-channel token-aligned interface through a gated residual pathway. GraphMessage applied a separate gated residual projection to the mean representations of incoming graph neighbours. In graph-on mode, the model received sequence-aligned features and within-window graph links. In graph-shuffle mode, feature rows were permuted within each example and destination indices were permuted after batch flattening. This retained the edge count and the marginal source out-degree and destination in-degree counts, while allowing cross-example links and changing source–destination pairings and local graph topology. Graph-off bypassed both pathways. The model contained 479, 195, 678 unique parameters (approximately 148.6 million active per token).

Pretraining combined masked language modelling with two graph-feature reconstruction loss terms: one over all feature-valid positions and one over masked-token positions. Both terms used the shared eight-dimensional graph-reconstruction head and were weighted at 0.25 relative to masked language modelling; MoE regularisation was also applied. Variant-type and contrastive path-context objectives were assigned zero weight and did not contribute to training. Gaussian noise (s.d. 1.0) was added to the graph-feature inputs, after which graph features at masked positions were set to zero before reconstruction. A five-stage curriculum increased model length from 512 to 8, 192 positions over 116, 883 optimisation steps. AdamW used a peak learning rate of 3 × 10−4, cosine decay to 1 × 10−5, 12, 000 warm-up steps, weight decay of 0.01 and gradient clipping at 1.0. Training used bfloat16 on four 80-GB NVIDIA A100 GPUs with an effective batch size of 64. The graph-sensitivity analysis was conducted under this combined production objective and was interpreted as a joint response to the supplied graph inputs, rather than as an attribution of an individual feature group, topology component or auxiliary loss.

### Graph-input sensitivity

At each stage-end checkpoint, 25, 000 training-panel windows were evaluated in graph-on, graph-shuffle and graph-off modes using identical batches of eight windows. Paired batch-level masked-language-model loss differences were summarised using 95% intervals from 1, 000 paired percentile-bootstrap resamples (Efron and Tibshirani, 1993), with Benjamini-Hochberg adjustment for predefined comparisons (Benjamini and Hochberg, 1995).

### Held-out sequence classification and adaptation

Gene-versus-intergenic and coding-sequence (CDS)-versus-intergenic datasets were generated separately for LA1974 and MicroTom using 512-bp windows. Positive windows were centred within annotated intervals, negative windows were entirely intergenic, and boundary-crossing windows were excluded. Classes were balanced. Chromosomes 03 and 05 were reserved for final testing, while all other chromosomes were used for fitting and validation.

Frozen representations were extracted from TomatoPGFM, DNABERT-2 (Zhou et al., 2024) and PlantDNAMamba (Liu et al., 2025). TomatoPGFM was evaluated without graph input and mean-pooled over its 87 non-padding positions, whereas baseline models used native tokenisers and attention-mask-aware mean pooling. For each model and task, StandardScaler and class-balanced logistic regression were fitted as a pipeline within each of three stratified training folds. The regularisation parameter C was selected from seven prespecified values by mean validation AUROC. The selected pipeline was refitted on all training-chromosome samples and evaluated once on chromosomes 03 and 05. AUROC was the primary metric, with 95% intervals estimated from 1, 000 class-stratified bootstrap resamples; secondary metrics were also calculated. Pooling sensitivity compared the 87-position mean, all 512 executed positions and direct unpadded execution. A zero-feature GraphAdapter control supplied all-zero graph features and no edges.

LoRA adaptation (Hu et al., 2022) used the same partitions and sequence-only TomatoPGFM input, with rank 8, alpha 16 and dropout 0.05. For TomatoPGFM, LoRA was applied to selected projection layers within the sequence backbone, while corresponding architecture-specific projection layers were adapted in the baseline models. Adapter and classifier learning rates were 2 × 10−4 and 1 × 10−3, with weight decay 0.01 and batch size 32.

Training ran for up to 15 epochs with patience 3; a random 10% subset of the training-chromosome samples was used for validation, and only adapters and the linear classifier were trainable. Gene test sets matched the frozen-probe analysis, and CDS test sets contained 767 LA1974 and 885 MicroTom windows.

### Inference benchmarking and statistical analysis

Inference was benchmarked on one 80-GB NVIDIA A100-SXM4 GPU in FP32 with batch size 16, three warm-up batches and 10 timed batches, and latency was measured over 30 repetitions. TomatoPGFM was tested with the zero-feature GraphAdapter pathway enabled and in graph-off mode at 512, 1, 024 and 2, 048 positions. The adapter-on condition used zero-valued graph features and edge_index = None, so adjacency-based GraphMessage aggregation was not executed. We recorded sequence throughput, median and 90th-percentile latency, and peak GPU memory.

## Data availability

The 66 training assemblies were obtained from Zhou et al. (2022; NCBI BioProject PRJNA733299), Li et al. (2023; NCBI BioProject PRJNA809001) and Shi et al. (2026; CNCB BioProject PRJCA030093, NCBI BioProject PRJNA1201608 and Zenodo https://doi.org/10.5281/zenodo.17878268). The LA1974 assembly is available under NCBI BioProject PRJNA633104 (Molitor et al., 2021). The MicroTom genome is available as NCBI RefSeq GCF_036512215.1 (O’Leary et al., 2016; Shirasawa and Ariizumi, 2024). Processed coordinate manifests, sample-level predictions and result tables generated in this study are available at Zenodo (https://doi.org/10.5281/zenodo.22033079) under CC BY 4.0. The original third-party assemblies and annotations are not redistributed by this study.

## Code and model availability

Source code is available at https://github.com/tomatoai2026/TomatoPGFM and is archived at https://doi.org/10.5281/zenodo.22035724 (concept DOI https://doi.org/10.5281/zenodo.22035723). The inference-only TomatoPGFM weights (model.safetensors; SHA-256 462f3bfc7178c4558fe993ef579d925a6a54a7448c975b8488bf74d0ac36f7c) are available from Hugging Face (https://huggingface.co/tomatoai2026/TomatoPGFM), ModelScope (https://modelscope.cn/models/turgun/TomatoPGFM) and Zenodo (https://doi.org/10.5281/zenodo.22032734). The optimizer-bearing training-resume checkpoint is not included in the public release.

## Author contributions

J. Han and T. Yushan contributed equally to this work. J. Han: Conceptualization, Methodology, Writing – original draft; T. Yushan: Model training, Software development, Writing – original draft; J. Wang, H. Yang, J. Zhao, F. Jiang, C. Jia and T. Yang: Data curation; B. Wang: Supervision, Project administration, Writing – review & editing; C. Zhang and Q. Yu: Conceptualization, Supervision, Project administration, Writing – review & editing. All authors reviewed and approved the final manuscript.

## Acknowledgements

The authors received no specific funding for this work and have no additional acknowledgements.

## Conflict of interest

The authors declare no competing interests.

## Supplementary Tables

**Table S1.** Composition and roles of the pretraining panel and held-out evaluation accessions.

| Group | Accessions | Sequence | Annotated genes | Role |
| --- | --- | --- | --- | --- |
| Training panel | 66 | 54.65 Gb | 2,269,697 | Graph construction and pretraining |
| LA1974 | 1 | 1,232.3 Mb | 36,233 | Held-out sequence-only evaluation |
| MicroTom | 1 | 832.8 Mb | 39,809 | Held-out sequence-only evaluation |

## Notes

https://github.com/tomatoai2026/TomatoPGFM

https://huggingface.co/tomatoai2026/TomatoPGFM

https://modelscope.cn/models/turgun/TomatoPGFM

https://doi.org/10.5281/zenodo.22033079

https://doi.org/10.5281/zenodo.22035724

https://doi.org/10.5281/zenodo.22032734

### Competing Interest Statement

The authors have declared no competing interest.

